# MACS: Rapid aqueous clearing system for three-dimensional mapping of intact organs

**DOI:** 10.1101/832733

**Authors:** Jingtan Zhu, Tingting Yu, Yusha Li, Jianyi Xu, Yisong Qi, Yingtao Yao, Yilin Ma, Peng Wan, Zhilong Chen, Xiangning Li, Hui Gong, Qingming Luo, Dan Zhu

## Abstract

Tissue optical clearing techniques have provided important tools for large-volume imaging. Aqueous-based clearing methods are known for good fluorescence preservation and scalable size maintenance, but are limited by either long incubation time, or insufficient clearing performance, or requirements for specialized devices. Additionally, due to the use of high concentration organic solvents or detergents, few clearing methods are compatible with lipophilic dyes while maintaining high clearing performance. To address these issues, we developed a rapid, highly efficient aqueous clearing method with robust compatibility, termed m-xylylenediamine (**M**XDA)-based **A**queous **C**learing **S**ystem (MACS). MACS can render intact organs highly transparent in a fairly short time and possesses ideal compatibility with multiple probes, especially for lipophilic dyes. Using MACS, we cleared the adult mouse brains within only 2.5 days for three-dimensional (3D) imaging of the neural structures labelled by various techniques. Combining MACS with DiI labelling, we visualized the vascular structures of various organs. MACS provides a useful tool for 3D mapping of intact tissues and is expected to facilitate morphological, physiological and pathological studies of various organs.

## Introduction

Three-dimensional (3D) imaging of tissue structures at high resolution plays an indispensable role in life science. The development of diverse fluorescent labelling methods and optical imaging techniques provides essential tools for 3D imaging of large-volume tissues^1–4^. However, the imaging depth is rather limited due to the opaqueness of tissue^5^. Automated serial-sectioning and imaging techniques have been developed to address this issue, allowing the acquisition of high-resolution images throughout the brain^6–9^.

As a distinct solution, tissue optical clearing technique has been proposed for imaging deeper without cutting^10–13^. In the past decade, a variety of optical clearing methods have been developed and are principally divided into solvent-based methods, such as 3DISCO^14, 15^, iDISCO^16^, uDISCO^17^, PEGASOS^18^, Ethanol-ECi^19^, FDISCO^20^, vDISCO^21^, sDISCO^22^, boneclear^23^ and aqueous-based methods, such as SeeDB/SeeDB2^24, 25^, *Clear^T^*^26^, Sca*l*eS^27^, CUBIC-series^28–31^, CLARITY^32, 33^, PACT-PARS^34^, SWITCH^35^, OPTIClear^36^ and Ce3D^37^. These methods provide powerful tools for visualizing tissue structures and greatly promote the development of life science.

Aqueous-based clearing methods are known for good fluorescence preservation and scalable maintenance of tissue size^11^, but are also faced with poor clearing performance or slow clearing speed. For example, SeeDB and *Clear^T^* performed well on embryos and neonatal mouse tissues, while showed modest transparency in whole-mount adult tissues^38, 39^. CUBIC and CLARITY can achieve high tissue transparency but require long incubation time for clarification (e.g., approximately 2-3 weeks for the whole brain). Moreover, due to the usage of organic solvents or high concentration detergents, most clearing methods with high clearing capability are incompatible with lipophilic dyes, such as DiI, which has been widely used to trace neuronal structures and vasculatures^26, 40–42^. These drawbacks have limited their applications in researches.

In this work, we developed an aqueous clearing method based on m-xylylenediamine (MXDA), termed **M**XDA-based **A**queous **C**learing **S**ystem (MACS). MACS achieved high transparency of intact organs and rodent bodies in a fairly short time and showed ideal compatibility with multiple probes, especially with lipophilic dyes. MACS is applicable for imaging the neural structures of transgenic whole adult brains and immunostained mouse embryos, as well as the neural projections throughout the whole brain labelled by viruses. MACS also allows imaging of DiI-labelled vascular structures of various organs. MACS is expected to promote comprehensive morphological and pathological studies of intact organs.

## Results

### MACS enables rapid clearing of multiscale tissues

We successfully cleared the intact brain with a high level of transparency within only 2.5 d using our MACS protocol (Fig. 1a). Compared with other available clearing methods, MACS renders brain samples highly transparent much faster (Fig. 1a-c, Fig. S1a, c) and nearly maintains the sample size after transient expansion (Fig. 1d, Fig. S1b). The computed tomography (CT) reconstruction images revealed no significant changes of brain volume before and after MACS clearing (Fig. S2a, b). The outlines and internal regions of the brain slices overlapped well before and after MACS treatment (Fig. S2c-h). Furthermore, to investigate the influence of MACS on the preservation of fine structures, we imaged a typical pyramidal neuron and single microglia before and after clearing. The results showed that MACS could maintain the cell morphology and fine structures well (Fig. S2i, j).

**Fig. 1.**
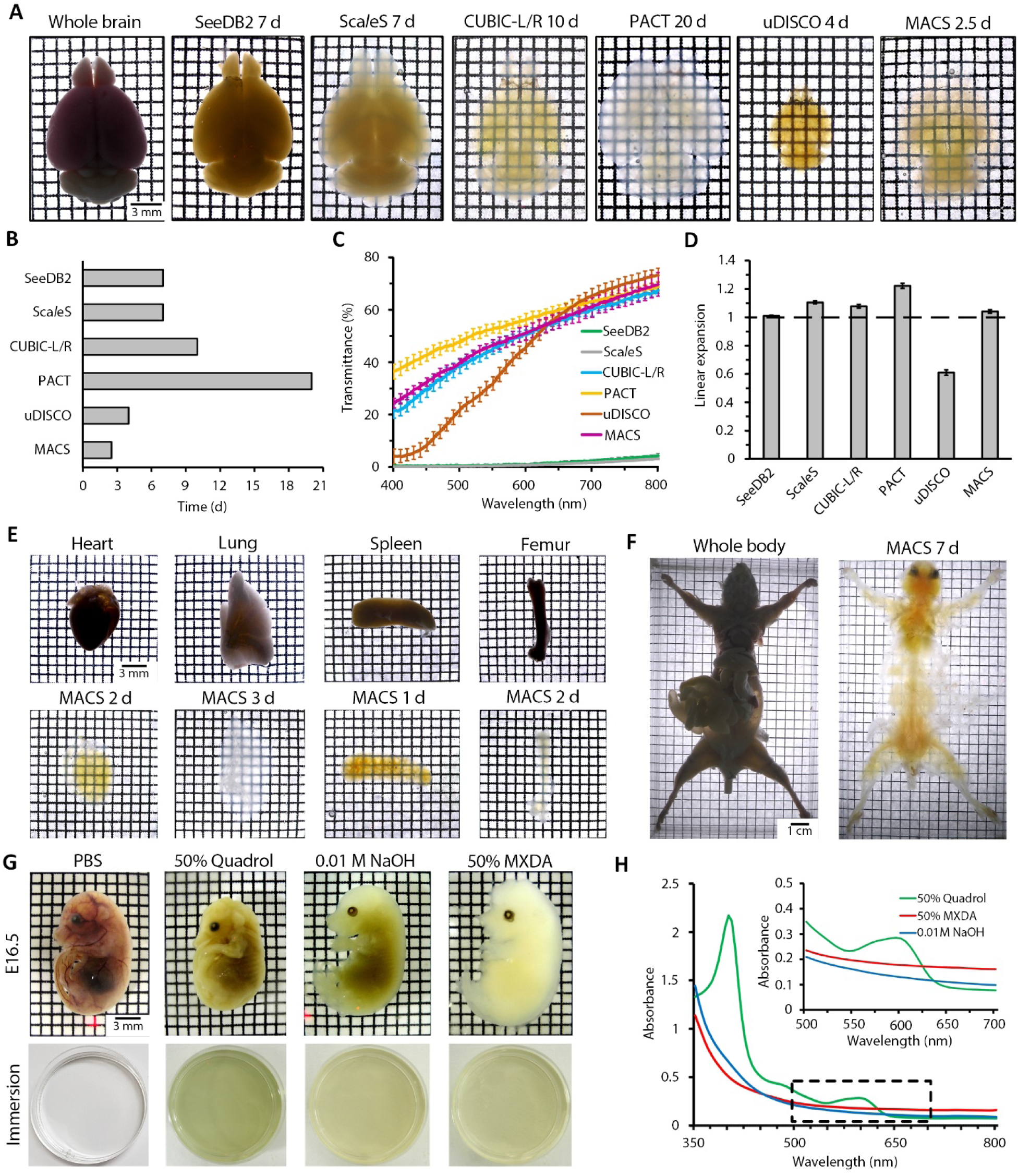
MACS enables rapid clearing for multiscale tissues with decolourization. (**a**) Bright field images of whole adult mouse brains cleared by different clearing protocols. (**b**) Comparison of clearing time needed for each method. (**c**) Transmission curves of cleared whole brains treated by different methods. Light transmittance was measured from 400 to 800 nm (n=3). (**d**) Quantification of linear expansion of whole brains after clearing by each method (n=3 brains each). (**e**) Clearing performance of both hard and soft organs cleared by MACS. (**f**) Clearing performance of MACS for P60 whole mouse body. (**g**) The decolourization effect of different solutions, including 50 vol% MXDA, 50 vol% Quadrol and 0.01 M NaOH. The reflection images of fixed embryos and immersions are shown before and after decolourization. Note that each immersion is colourless before decolourization. (**h**) The absorbance curves of each solution were measured after decolourization. Inset: magnification of the boxed region. All values are presented as the mean ± s.d.

MACS could also efficiently clarify other mouse tissues, including both soft internal organs and hard bones (Fig. 1e), and was also applicable for the adult mouse body (Fig. 1f). Additionally, MACS was effective for intact adult rat organs (Fig. S3a). Notably, MACS demonstrated superior ability in clearing mouse embryos and pups (Fig. S3b).

We also found that MXDA solution could efficiently decolourize heme-rich tissues, such as embryos (Fig. 1g). The absorbance of the decolourizing medium indicated that MXDA decolourized the samples in a manner similar to that of NaOH solution (release Fe) but quite different from that of Quadrol solution (release heme) used in CUBIC^29^ (Fig. 1h). Additionally, MXDA showed high pH stability over NaOH solution during clearing (Table S1). The decolourizing capability of MXDA enables MACS to decolourize samples during clarification (Fig. S3c).

### MACS preserves signals of multiple fluorescent probes, especially lipophilic dyes

Furthermore, we investigated the fluorescence preservation of MACS for both endogenous fluorescent proteins and chemical fluorescent tracers. The results demonstrated that MACS could preserve the endogenous EGFP, EYFP and tdTomato very well after clearing (Fig. 2a, b, Fig. S4a). We also imaged MACS-cleared brain slices during long-term storage, and the fluorescence intensity remained relatively high after one month (Fig. 2c, Fig. S4b).

**Fig. 2.**
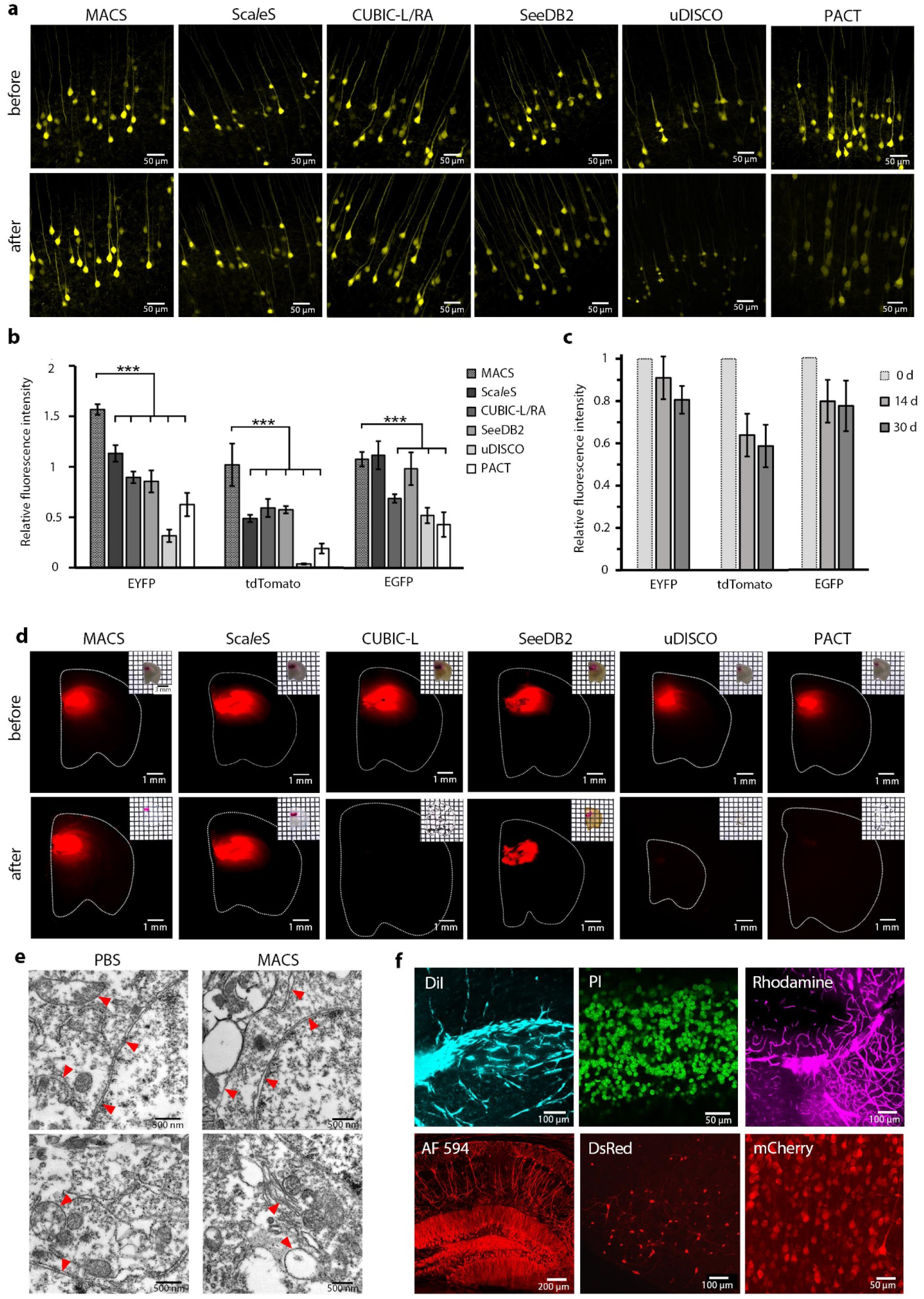
MACS maintains the signals of multiple fluorescent probes. (**a)** Fluorescence images of endogenous EYFP signals (1 mm *Thy1*-YFP-H brain slices) before and after MACS clearing compared with other clearing protocols. (**b**) Quantification of fluorescence preservation of EYFP, tdTomato and EGFP after MACS clearing compared with different methods (n=3). (**c**) Quantitative analysis of long-term fluorescence preservation of EYFP, tdTomato and EGFP after MACS clearing (n=3). (**d**) Bright field and fluorescence images of DiI-labelled brain slices before and after clearing by each method. (**e**) Ultramicroscopic imaging of mouse brain samples restored from MACS or PBS by transmission electron microscopy. Red arrow heads indicate typical membrane structures. (**f**) Fluorescence signals labelled by various chemical fluorescent tracers are finely imaged after MACS clearing, including DiI, PI, Tetramethylrhodamine (Rhodamine), AlexaFluor 594 (AF 594)-conjugated antibody, DsRed and mCherry. All values are presented as the mean ± s.d.; Statistical significance in **b** (***, *P* < 0.001) was assessed by one-way ANOVA followed by the Bonferroni post hoc test.

DiI is a commonly used lipophilic fluorescent dye for neural tracing and vascular labelling. Due to the high concentration of membrane-removing detergents or organic solvents, CUBIC-L, PACT and uDISCO are not compatible with DiI labelling. By contrast, MACS can preserve the fluorescence of DiI fairly well, similar to Sca*l*eS and SeeDB2 (Fig. 2d). Indeed, MACS used neither detergents nor organic solvents, thus samples treated by MACS maintained membrane integrity well, which was crucial for DiI signalling (Fig. 2e). The superior compatibility enables visualization of neuronal projections in the DiI-labelled hippocampus region in mouse brain tissue (Fig. 2f). We also tested MACS with other types of chemical tracers, including propidium iodide (PI) for nuclear staining, Tetramethylrhodamine-conjugated dextran for vessel labelling, virus-delivered proteins (DsRed and mCherry) and fluorophore-conjugated antibodies in immunostaining (AlexaFluor 594 (AF 594) and AlexaFluor 633 (AF 633)) (Fig. 2f, Fig. S4c, d). The results showed that MACS maintained the fluorescence signals of all tested tracers well.

Additionally, previous studies demonstrated that CM-DiI could be used as an alternative in CLARITY-based methods. CM-DiI is an aldehyde-fixably modified version of DiI which adheres not only to the cellular membranes but also protein structures after fixation, such that it would remain in the tissue after lipid extraction^43^. However, we found that the signals from CM-DiI was only partial remained after treatment by CUBIC-L, PACT and uDISCO, the signal loss was obvious as previously reported^44^. Due to the good membrane integrity after MACS treatment, both the DiI and CM-DiI signals were well maintained without any obvious loss (Fig. S4e).

### MACS achieves 3D reconstruction of neural structures in intact tissues

Using the rapid MACS clearing protocol, we cleared transgenic whole mouse brains and performed fluorescence imaging by light sheet fluorescence microscopy (LSFM). We obtained the neurons in *Thy1*-GFP-M mouse brain at different depths and performed 3D reconstruction (Fig. 3a-d). The fine neural structures were well observed in different brain regions, including the midbrain, hippocampus, cerebellum, striatum, cortex, and cerebellar nuclei (Fig. 3e-j). We also cleared and imaged the *Sst*-IRES-Cre::Ai14 transgenic mouse brain, and MACS allowed fine imaging of tdTomato fluorescence labelled neurons throughout the whole brain at single-cell resolution (Fig. S5a-e).

**Fig. 3.**
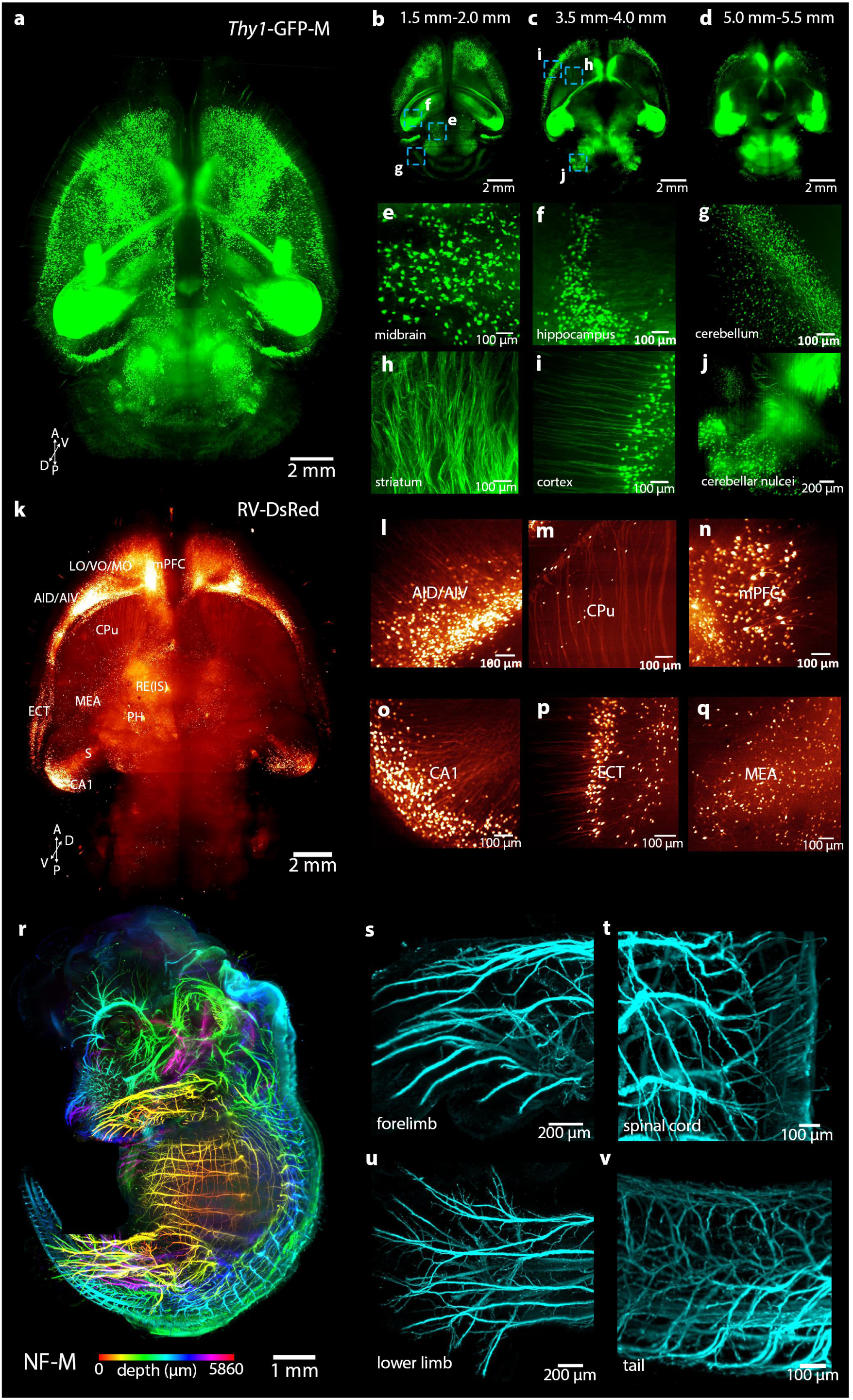
MACS is applicable for 3D imaging and reconstruction of neural structures in intact tissues. (**a**) 3D reconstruction of LSFM images of whole brain (*Thy1*-GFP-M) cleared by MACS. (**b-d**) Maximum projection of acquired images between 1.5 mm-2.0 mm (**b**), 3.5 mm-4.0 mm (**c**) and 5.0 mm-5.5 mm (**d**). (**e-j**) The high-magnification images of cleared brains reveal fine structures at different positions of the brain, including the midbrain (**e**), hippocampus (**f**), cerebellum (**g**), striatum (**h**), cortex (**i**), and cerebellar nuclei (**j**). (**k**) 3D reconstruction of RV-labelled afferent projections to nucleus reuniens (RE) throughout the whole brain. (**l-q**) Several regions of specific projections to RE, including the dorsal/ventral agranular insular cortex (AID/AIV) (**l**), caudate putamen (CPu) (**m**), medial prefrontal cortex (mPFC) (**n**), ventral CA1 of the hippocampus region (**o**), ectorhinal cortex (ECT) (**p**) and medial amygdaloid nucleus (MEA) (**q**). (**r**) 3D reconstruction of whole embryo (E14.5) labelled for neurofilament (NF-M). The images along the z stack are coloured by spectrum. (**s-v**) Details of the nerve innervation in the forelimb (**s**), spinal cord (**t**), lower limb (**u**), and tail (**v**). Thin nerve fibres are finely labelled and detected.

Virus labelling is widely used to reveal neural circuits across the whole brain^45, 46^. Here, we applied MACS protocol to visualize neuronal projections labelled by different types of virus, including retrograde rabies virus (RV) and anterograde adeno-associated virus (AAV). We imaged the input and output of the nucleus reuniens (RE), which reportedly receives afferent projections across the brain and is an important temporal constraint in hippocampus-RE-mPFC circuits^47, 48^. We injected RV-DsRed and AAV-mCherry into the RE region for retrograde and anterograde tracing of the projections, respectively (Fig. S5f). The 3D rendering of acquired images showed a diverse and widely distributed set of afferents to RE (Fig. 3k), which heavily projected from the dorsal/ventral agranular insular cortex (AID/AIV), medial prefrontal cortex (mPFC), ventral CA1 of the hippocampus, ectorhinal cortex (ECT), medial amygdaloid nucleus (MEA) (Fig. 3l-q), etc. Moreover, the 3D rendering showed a relatively limited output from RE, which was mainly directed to the hippocampal formation (e.g., CA1), ventral subiculum (S) and mPFC (Fig. S5g-k), as previously reported^49^.

In addition to transgenic labelling and virus labelling, immunostaining is a powerful method to label tissues. Due to the hyperhydration of MXDA used in MACS, we explored whether MXDA pretreatment would enhance the permeability of antibodies in immunostaining. The results revealed that MXDA pretreated samples could achieve deeper staining than those without MXDA pretreatment (Fig. S6a-c). After MXDA pretreatment, whole embryos were stained with neurofilament antibody and followed by MACS clearing and imaging with LSFM (Fig. S6d). We obtained the 3D nerve distributions of embryos at different ages (Fig. 3r, Fig. S6e, j). The fine neural branches in the limbs, spinal cord, tail, and whisker pad could be clearly visualized (Fig. 3s-v, Fig. S6f-i).

### MACS enables 3D mapping of the DiI-labelled vascular system

High-resolution reconstruction of the vasculature of various organs facilitates studies of many vascular-associated diseases^50, 51^. A previous study provided an excellent labelling protocol for vasculatures with high signal intensities by perfusion of DiI solution^52^, which has been demonstrated experimentally to be more effective on specific mouse organs (e.g. mouse spleen and kidney) than some common-used labelling methods (Fig. S7). However, few clearing methods could be applied to this labelling because of incompatibility. Here, due to the superior compatibility with DiI and high clearing capability, we applied MACS to acquire the vasculatures of DiI-labelled organs, including the whole mouse brain, spinal cord and other internal organs.

After MACS clearing, the DiI-labelled vasculature of the whole brain could be observed directly (Fig. 4a). Combined with LSFM imaging, we visualized the brain vasculature in 3D (Fig. 4b). The detailed vascular structures in the cortex, middle of the brain, cerebellum and hippocampus could be clearly identified (Fig. 4c-f). The sagittal view showed the vascular distribution along the z-axis (Fig. 4g). The spinal cord was also finely imaged with both the central blood vessels and the surrounding capillaries distinguished (Fig. 4h, i). The vascular network throughout the entire spleen was constructed with fine vessels clearly visualized (Fig. 4j, k)

**Fig. 4.**
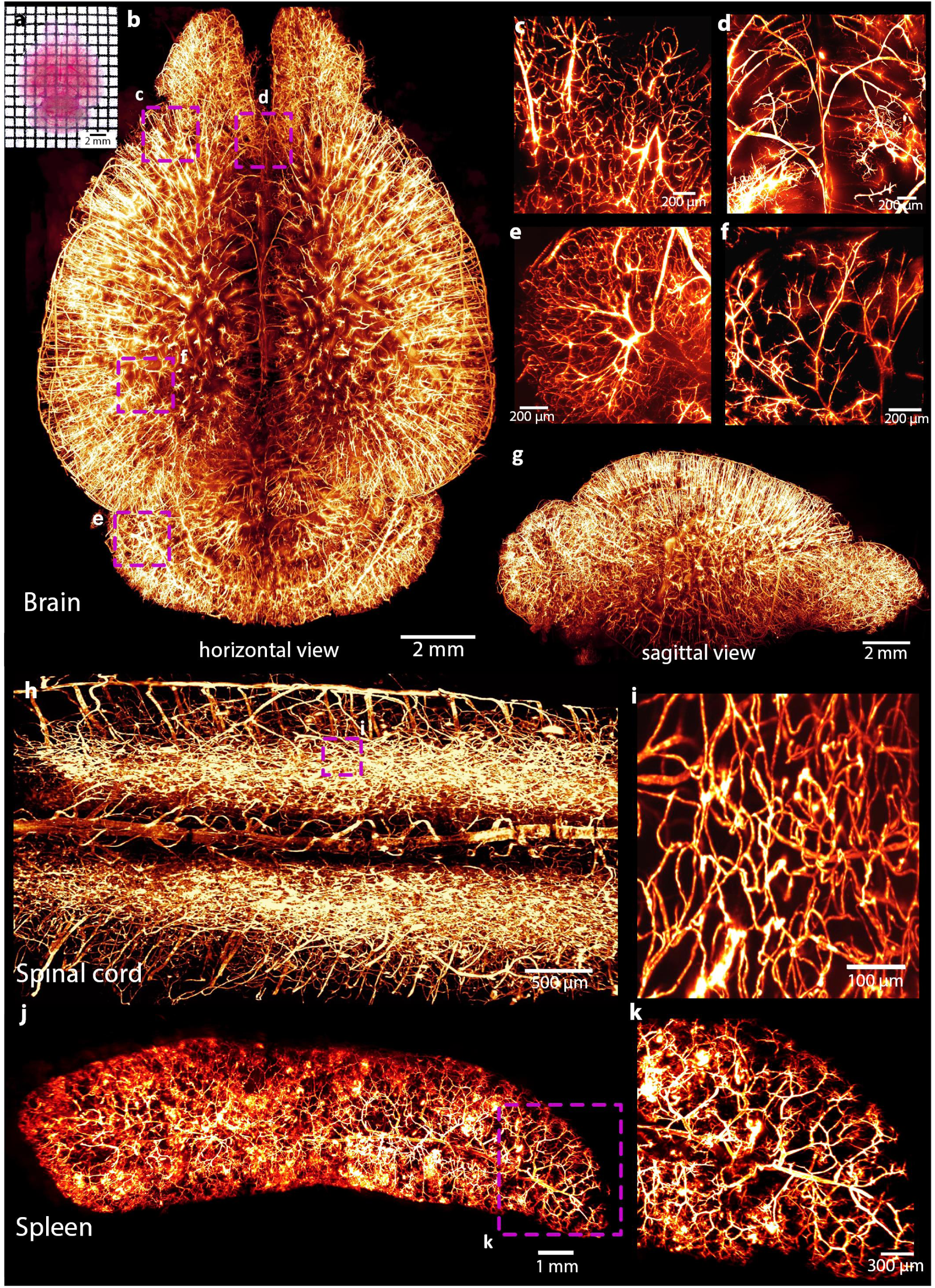
MACS enables 3D visualization of the vascular network of DiI-labelled mouse organs. (**a**) Direct view of DiI-labelled vasculatures in the cleared brain. (**b**) 3D rendering of the vascular network throughout an entire adult brain imaged by LSFM. (**c-f**) Detailed vasculature in the cortex (**c**), middle of the brain (**d**), cerebellum (**e**) and hippocampus (**f**). (**g**) Sagittal view of the reconstructed brain shown in **a**. (**h**) 3D rendering of vasculatures in the spinal cord. (**i**) Magnification of boxed regions in **h**, the surrounded capillaries are finely labelled and clearly visualized. (**j**) Reconstructed vascular network of adult mouse spleen. (**k**) Magnification of boxed regions in **j**, the small branches are well detected.

## Discussion

For current aqueous-based clearing methods, the main urgent issue is that methods with good clearing capability usually need rather long incubation time or complex equipment for clearing, and methods with reduced clearing time always show insufficient transparency on adult organs. In this article, we solved this problem by presenting MACS, a rapid and effective aqueous-based optical clearing method scalable for various tissues. MACS can make samples highly transparent in a fairly short time. For instance, MACS requires only 2.5 d to clear a whole adult brain, saving almost 80% of the time needed for CUBIC. MACS also shows ideal compatibility with multiple probes, especially with lipophilic dyes. Combined with LSFM, MACS performed well in 3D reconstruction of neuronal structures in various intact tissues. Due to its superior compatibility with DiI, MACS is applicable to 3D visualization of DiI-labelled vascular structures of various tissues.

Notably, MXDA was first introduced to tissue clearing and used as the central reagent in MACS. Similar to urea, MXDA has two NH2 groups, which indicates a strong hydration ability to release dense fibres. Moreover, MXDA has a rather high RI up to 1.57. Combined with its superior liquidity and water solubility, MXDA can efficiently decrease the RI mismatch in tissue and contribute to the rapid clearing capability of MACS, addressing the problem of long clearing time (weeks to months) of the available aqueous methods. The addition of sorbitol further enhances transparency and fluorescence preservation. By controlling the concentration of MXDA and sorbitol, the MACS protocol was designed for three steps by passive immersion, which is rather rapid, highly efficient and simply executed.

Recently, several groups have introduced novel physical principles and devices to accelerate the CLARITY clearing procedure, such as stochastic electrotransport^53^ and ACT-PRESTO^54^. The acceleration is very successful that whole brains could be clarified within several days, but requires highly specialized devices. For MACS, a tri-step incubation with no need of extra equipment makes it rather simple and convenient for researchers to achieve rapid and high-performance clearing using this method. Additionally, the solvent-based clearing methods can also achieve high transparency in a relatively short time but often exhibit fluorescence quenching and tissue shrinkage. Recently, the problem of fluorescence quenching has been partially resolved by many groups^17, 18, 20^. However, the tissue shrinkage caused by thorough dehydration is inevitable. MACS could not only nearly maintain the original sample size with tissue fine structures preserved but also achieve high transparency in a fairly short time, it may become beneficial for such studies.

For some heme-rich tissues, such as the spleen, kidney and embryo, residual blood in these tissues will cause severe light absorbance in the visible region (400-600 nm)^55^, thus affecting tissue transparency and imaging quality. To overcome this limitation, several effective chemicals showing good decolourizing capability have been used for decolourization in the latest clearing methods^18, 21, 29, 31^. In this study, we found that MXDA also had excellent decolourizing characteristic. The decolourization principle of MXDA is different from that of Quadrol (releasing hemin) used in CUBIC but similar to that of NaOH (releasing Fe)^29^. However, during immersion, the pH of NaOH solution showed an obvious decrease, while MXDA showed high pH stability (Table S1). The decolourization effect of MXDA enables MACS to directly extract residual chromophores in tissues, thus leading to efficient clearing without any additional decolourization steps. Furthermore, MACS is expected to be combined with solvent-based methods, which are often blood-sensitive, to provide decolourization prior to clearing.

Compatibility with diverse fluorescent labels is also essential for a successful clearing method. MACS shows great compatibility with both endogenous fluorescence proteins and many chemical fluorescent tracers, and is also compatible with virus labelling and immunostaining. We also find that pretreatment with MXDA solution could enhance the permeability of tissues and promote antibody penetration. Moreover, the immunostained samples nearly maintained their original sizes after MACS clearing and could be finely imaged by LSFM. This method will provide valuable tool in studies that require whole-mount immunostaining and imaging for large tissues with minimal size changes.

Lipophilic carbocyanine dyes, such as DiI, have been widely used to label cell membranes and trace neuronal projections in live and fixed tissues^26, 40–42^. These dyes can also be used to attain fine labelling of the vasculature^52^. However, existing methods with high clearing capability are always incompatible with lipophilic dyes due to the use of high concentrations of detergents or organic solvents. MACS uses neither solvents nor detergents during clearing, so it preserves the membrane architectures that are critical for DiI labelling thus demonstrates ideal compatibility with lipophilic dyes (Fig. 2d, e), along with its high clearing performance, MACS enables 3D visualization of DiI-labelled vascular structures in various intact organs (Fig. 4). Notably, the DiI labelling method could offer more detailed vascular information than other commonly used methods in specific organs. This approach is expected to facilitate the analysis of vasculature networks in specific disease models. Recently, CM-DiI has been reported to be used as an alternative of DiI in CLARITY method^43^. However, CM-DiI was experimentally only partial compatible with methods using high concentration of detergents or organic solvents, such as CUBIC-L, PACT and uDISCO. The signal loss was obvious, which was consistent with previously reported^44^. For MACS, both the DiI and CM-DiI signals were well maintained without obvious loss. Additionally, the CM-DiI dye is nearly 100 times more expensive than the common DiI dye (price from Thermo Fisher). It seems not cost-effective to label the vasculature by perfusing large amount CM-DiI dyes. So we believe the excellent DiI compatibility along with the high clearing performance is a big advantage for MACS over existing methods.

For aqueous-based methods, high RI aqueous solutions are often used for final matching with scatters such as lipid or protein^11^. High concentration sugars and polyalcohols are often employed in matching solution. However, most of these solutions have limited RI and high viscosity. Contrast reagents, such as iohexol solution, can achieve high RI, but too costly for common use. In MACS, due to the high RI of MXDA (up to 1.57), the MACS-R2 has a high RI of 1.51 with low viscosity, which was sufficient for rapid RI matching and LSFM imaging. Additionally, the fluorescence signals are well maintained over time in MACS-R2 (Fig. 2c). In fact, the RI of MACS-R2 could be easily increased by adding extra sorbitol, which could be used for other experimental purpose, such as imaging with oil-immersion objectives (RI≈1.52). Notably, the price of MACS ingredient is rather cheap (about 1.3 US dollars per 10 ml, Tokyo Chemical Industry) for researchers to afford. These findings implies that the MACS-R2 is hopeful to be used widely as RI matching solution or mounting media in different studies.

As a newly developed clearing protocol, MACS not only maintains the common advantages of aqueous-based methods, including good fluorescence preservation and fine size maintenance, but also overcomes the limitations of slow clearing speed and insufficient transparency. MACS is also compatible with lipophilic fluorescent dyes. In summary, MACS is a rapid, highly efficient clearing method with robust compatibility. We believe that MACS could provide a valuable alternative for the clearing, labelling and imaging of large-volume tissues. Moreover, MACS has a great potential for 3D pathology of human clinical samples.

## Materials and Methods

### Animals

Animals were housed in a specific-pathogen-free (SPF) animal house under a 12/12 h light/dark cycle with unrestricted access to food and water. Wild-type (C57BL/6J, 8-12 weeks old), *Thy1*-GFP-M (8-10 weeks old), *Thy1*-YFP-H (8-10 weeks old), *Sst-IRES-Cre*::Ai14 (8-9 weeks old), *Cx3cr1*-GFP (8-12 weeks old) mice and Sprague-Dawley rats (8 weeks old) were used in this study. Animals were selected for each experiment based on their genetic background (wild-type or fluorescence transgenes). All animal experiments protocols were performed under the Experimental Animal Management Ordinance of Hubei Province, P. R. China, and the guidelines from the Huazhong University of Science and Technology. These protocols were approved by the Institutional Animal Ethics Committee of Huazhong University of Science and Technology.

### Sample preparation

Adult mice and rats were deeply anaesthetized with a mixture of 2% α-chloralose and 10% urethane (8 ml/kg) and were transcardially perfused with 0.01 M phosphate buffered saline (PBS, Sigma, P3813) followed by 4% paraformaldehyde (PFA, Sigma-Aldrich, 158127) in PBS for fixation. The intact brains, bones, and desired organs were excised from the perfused animal body. Mouse embryos were collected from anaesthetized pregnant mice. The day on which a plug was found was defined as embryonic day 0.5 (E0.5). All harvested samples were post-fixed overnight in 4% PFA at 4°C. Coronal brain sections (1 mm and 2 mm) were sliced using a commercial vibratome (Leica VT 1200 s, Germany).

### Preparation of MACS solutions

The MACS protocol involves three solutions, which were consisted of gradient MXDA and sorbitol dissolved in distilled water or PBS, termed MACS-R0, MACS-R1 and MACS-R2, respectively. Proper heating with a water bath could promote the dissolution of sorbitol. When preparing MACS solutions, latex gloves should be wore to avoid direct contact with chemicals.

### MACS clearing procedure

For passive clearing, fixed samples were serially incubated in 20-30 ml MACS-R0, MACS-R1 and MACS-R2 solutions in 50 ml conical tubes with gentle shaking. The time needed for clearing in each solution depends on the tissue type and thickness. Hard bones and whole body should be incubated first in 0.2 M ethylene diamine tetraacetic acid (EDTA) (Sinopharm Chemical Reagent Co., Ltd, China) for decalcification and then treated with MACS-R0, MACS-R1 and MACS-R2 in succession. The incubation time in final buffer was adjustable by visual inspection for desired transparency. All other clearing protocols mentioned in this paper, including SeeDB2, Sca*l*eS, CUBIC-L/R/RA, PACT and uDISCO, were performed following the original publications^17, 24, 27, 31, 34^.

### Imaging

Fluorescence images of cleared samples (embryos from E12.5 to E16.5, whole adult brains and entire organs) were acquired with a light sheet microscope (Ultramicroscope I, LaVision BioTec, Germany), which was equipped with a sCMOS camera (Andor Neo 5.5) and a macrozoom body (Olympus MVX-ZB10, magnification from 0.63× to 6.3×) with a 2× objective lens (Olympus MVPLAPO2X, NA = 0.5, working distance (WD) = 20 mm). Thin light sheets were illuminated from both the right and left sides of the sample, and a merged image was saved. We acquired light-sheet microscope stacks using ImSpector (Version 4.0.360, LaVision BioTec) as 16-bit grayscale TIFF images for each channel separately.

An inverted laser-scanning confocal fluorescence microscope (LSM710, Zeiss, Germany) was used to perform fluorescence imaging of regions of brain sections and intact bones. Samples were placed on a slide with MACS-R2 and covered with a coverslip to keep tissue submerged in solutions. A 5× objective lens (FLUAR, NA=0.25, WD=12.5 mm), 10× objective lens (FLUAR, NA=0.5, WD=2 mm), and a 20× objective lens (PLAN-APOCHROMAT, NA=0.8, WD=550 μm) were used for imaging. Zen 2011 SP2 (Version 8.0.0.273, Carl Zeiss GmbH) software was used to collect data.

Specimens including brain slices were immersed in PBS or MACS-R2 and imaged before or after clearing with a fluorescence stereomicroscope (Axio Zoom. V16, Zeiss, Germany) using a 1× long working distance air objective lens (PLAN Z 1×, NA=0.25, WD = 56 mm). ZEN 2012 (Version 1.1.2.0, Carl Zeiss GmbH) was used to collect data.

### Transmission Electron Microscopy

Fixed brain slices were sequentially treated in MACS solution for 6-8 h each step, or stored in PBS at 4°C. The treated samples were then restored by washing in 1× PBS at room temperature for 12 h. The samples were post-fixed with 2.5% glutaraldehyde for several hours, then washed by 0.1 M PBS three times. The samples were then fixed with 1% Osmic acid in 0.1 M PBS buffer for 2 hours at room temperature, and washed three times in 0.1 M PBS buffer. The fixed samples were then dehydrated with a series of 30%, 50%, 70%, 90%, 100% ethanol. A 1:2 and 1:1 mixtures of epoxy resin and acetone were then sequentially added to infiltrate the blocks for 8-12 h at 37°C. The blocks were then incubated in pure epoxy resin at 37°C overnight, and allowed to polymerize at 65°C for 2 d. Thin sections were obtained using a Leica EM UC7 Ultramicrotomy. Sections were stained with uranyl acetate and lead acetate, and visualized using a Transmission Electron Microscope (FEI Tecnai G^2^ 20 TWIN, USA).

### Measurement of light transmittance

We measured the light transmittance of 2 mm thick mouse brain sections with a commercially available spectrophotometer (Lambda 950, PerkinElmer, USA). Cleared samples were placed on two glass slides covered with black tape, and a customized 3 mm× 3 mm slit was opened to obtain the collimated transmitted beam. We measured transmittance spectra from 400 nm to 800 nm.

### Measurement of tissue volume

To measure the volume of the whole mouse brain, a customized microcomputed Tomography (micro-CT) was used^56^. The whole mouse brains were imaged by micro-CT before and after MACS clearing, and reconstructed in 3D. The volume was calculated by Imaris software.

### Vasculature labelling

DiI stock solution was prepared by dissolving 30 mg DiI powder (Aladdin, D131225) in 10 ml 100% ethanol and stored in the dark at room temperature. DiI working solution was prepared by adding 200 ml DiI stock solution into 10 ml diluent (0.01 M PBS and 5% (wt/vol) glucose at a ratio of 1:4). DiI working solution should be freshly made under room lighting. CM-DiI (Thermo Fisher Scientifc, USA) solution was prepared as 0.01% (wt/vol). Anaesthetized mice were first perfused with 0.01 M PBS at a rate of 1-2 ml/min (total 3-4 min) and perfused with 10-15 ml DiI working solution (CM-DiI solution) at a rate of 1-2 ml/min (total 10-15 min). During perfusion with DiI solution, the ears, nose and palms will turn slightly purple. Finally, the mice were perfused with 4% PFA for fixation. Tissues of interest could be harvested and post-fixed in 4% PFA overnight.

Tetramethylrhodamine dextran was used to label the vasculature. Dextran (70,000 MW, Lysine Fixable, Invitrogen) was diluted in saline at a concentration of 15 mg/ml and injected into the tail vein (0.1 ml per animal). After injection, the animals were placed in a warm cage for 15-20 min before standard perfusion steps. Notably, heparin should not be added to PBS used in the prewashing step of the perfusion (use 0.1 M PBS instead), which will result in better labelling of the vasculature.

DyLight 649 conjugated L. esculentum (Tomato) lectin (LEL-Dylight649, DL-1178, Vector Laboratories) and Alexa Fluor 647 conjugated anti-mouse CD31 antibody (CD31-A647, 102416, BioLegend) were also used to label the vasculature. Lectin were diluted in saline to a concentration of 0.5 mg/ml and injected into the tail vein (0.1 ml per mouse). The Alexa Fluor 647 anti-mouse CD31 antibody (10 to 15 mg) was diluted in saline and injected into the tail vein (total volume of 200 ml). After injection, the animals were placed in a warm cage for 30 min prior to perfusion.

### Immunostaining

The following primary antibodies were used in this study: anti-GFP (Abcam, Ab290, dilution 1:500), anti-neurofilament (NF-M) (DSHB, 2H3, dilution 1:100), anti-beta-tubulin (Servicebio, GB13017-2, dilution 1:500), and anti-tyrosine hydroxylase (Servicebio, GB11181, dilution 1:500). Secondary antibodies including Alexa Fluor 594 goat anti-rabbit IgG (H+L) (1:500 dilution; A-11037, Life Technologies) and Alexa Fluor 633 goat anti-rabbit IgG (H+L) (1:500 dilution; A-21070, Life Technologies) were used.

For immunostaining without pretreatment, fixed brain slices were directly subjected to immunostaining with the primary antibodies in 1-2 ml 0.1% PBST (0.1 vol% Triton X-100 in PBS) containing 0.5% (w/v) donkey serum albumin and 0.01% (w/v) sodium azide for 2-3 d at room temperature with rotation. The stained samples were then washed with 10 ml 0.1% PBST several times with rotation and immersed in secondary antibodies in 1-2 ml 0.1% PBST containing 0.1% (w/v) donkey serum albumin and 0.01% (w/v) sodium azide for 2-3 d at room temperature with rotation. The samples were then washed with 10 ml 0.1% PBST several times and stored in PBS at 4°C prior to clearing.

For pretreatment, thick samples, such as brain slices, were stained using the following protocol: fixed samples were serially treated with MXDA solutions for 1-2 d and then washed with PBS several times. The recovered samples were subjected to immunostaining with primary antibodies in 1-2 ml 0.1% PBST (0.1 vol% Triton X-100 in PBS) containing 0.5% (w/v) donkey serum albumin and 0.01% (w/v) sodium azide for 2-3 d at room temperature with rotation. The stained samples were then washed with 10 ml 0.1% PBST several times with rotation and immersed in secondary antibodies in 1-2 ml 0.1% PBST containing 0.1% (w/v) donkey serum albumin and 0.01% (w/v) sodium azide for 2-3 d at room temperature with rotation. The samples were then washed with 10 ml 0.1% PBST several times and stored in PBS at 4°C prior to clearing.

For samples with large volumes, such as whole embryos, we used the following iDISCO immuno protocols. Fixed samples were serially treated with MXDA solutions for 1-2 d and then washed with PBS several times. The recovered samples were subsequently transferred to 50% methanol in PBS for 1 h, 80% methanol for 1 h, and 100% methanol for 1 h twice. Samples were then bleached with 5% H2O2 in 20% DMSO/methanol (vol%) at 4°C overnight. After bleaching, samples were washed with 100% methanol for 1 h twice, 80% and 50% methanol for 1 h, PBS for 1 h twice, and finally 0.2% PBST (0.2 vol% Triton X-100 in PBS) for 1 h twice. For the immunostaining step, pretreated samples were incubated in 0.2% PBST containing 20% DMSO and 0.3 M glycine at 37°C overnight, blocked in 0.2% PBST containing 10% DMSO and 6% goat serum at 37°C for 1 d, washed in PTwH (PBS–0.2% Tween-20 with 10 mg/ml heparin) overnight and then incubated with primary antibody dilutions in PTwH containing 5% DMSO and 3% goat serum at 37°C with gentle shaking on a shaker for 5-7 d. Samples were then washed for 2 d with PTwH and then incubated with secondary antibodies diluted in PTwH containing 3% goat serum at 37°C with gentle shaking for 5-7 d. Sections were finally washed in PTwH for 2 d and stored in PBS at 4°C prior to clearing.

### Neuronal tracing by RV and AAV

In this study, we used RV-N2C (G)-ΔG-DsRed (RV-DsRed, BrainVTA, R03002), rAAV-hSyn-mCherry-WPRE-pA (AAV-mCherry, BrainVTA, PT-0100) for tracing neuronal projections. For the RV and AAV injection to RE, 2-month-old C57BL/6J mice were used. The injection site of RV-DsRed (400 nl) and AAV-mCherry (400 nl) was targeted to the RE in different mice with the following coordinates: bregma, −0.82 mm; lateral, −0.27 mm; and ventral, 1.75 mm.

For injection, a cranial window was created on the skull to expose the brain area targeted for tracing neurons. The virus was injected into the brain using a custom-established injector fixed with a pulled glass pipette. The animal was placed in a warm cage after injection for waking up and then transferred into a regular animal room. The animals were kept for 7 d after RV and 28 d after AAV injection before perfusion.

### Image data processing

All raw image data were collected in a lossless TIFF format (8-bit images for confocal microscopy and 16-bit images for light sheet microscopy). Processing and 3D rendering were executed by a Dell workstation with 8 core Xeon processor, 256 GB RAM, and Nvidia Quadro P2000 graphics card. We used Imaris (Version 7.6, Bitplane AG) and Fiji (Version 1.51n) for 3D and 2D image visualization, respectively. Stitching of tile scans from light sheet microscopy was performed via Matlab (Version 2014a, Mathworks). The 16-bit images were transformed to 8-bit images with Fiji to enable fast processing using different software.

### Quantifications

#### Measurement of linear size changes

For the measurement of sample expansion and shrinkage, 2 mm brain slices and whole brains were used, and bright field images were taken before and after clearing. Based on top view photos, the area of samples was determined using the ‘polygon-selections’ function of Fiji. The linear expansion value was determined by the square root of the area size changes.

#### Relative fluorescence quantification

For evaluation of relative fluorescence intensity, the cell body of a neuron was encircled by the ‘freehand-selection’ tool in Fiji, and the mean fluorescence intensity and area were measured. The multiplication values of the two parameters were identified as the total fluorescence intensity of the neurons. The total fluorescence intensity was normalized to intensity in PBS (100%) for the same neuron, which was defined as relative fluorescence.

### Statistical analysis

Data are presented as the mean ± s.d. and were analysed using SPSS software (Version 22, IBM, USA) with 95% confidence interval. Sample sizes are indicated in the figure legends. For analysis of statistical significance, the normality of the data distribution in each experiment was checked using the Shapiro-Wilk test. The variance homogeneity for each group was evaluated by Levene’s test. P values were calculated using an independent-sample t test (two-sided) to compare data between two groups in Supplementary Information, Fig. S4b. P values were calculated using one-way ANOVA followed by the Bonferroni post hoc test to compare data in Fig. 2b, Supplementary Information and Fig. S1g. In this study, P < 0.05 was considered significant (*, P < 0.05; **, P < 0.01; ***, P < 0.001).

## Supplementary Figures

**Fig. S1.**
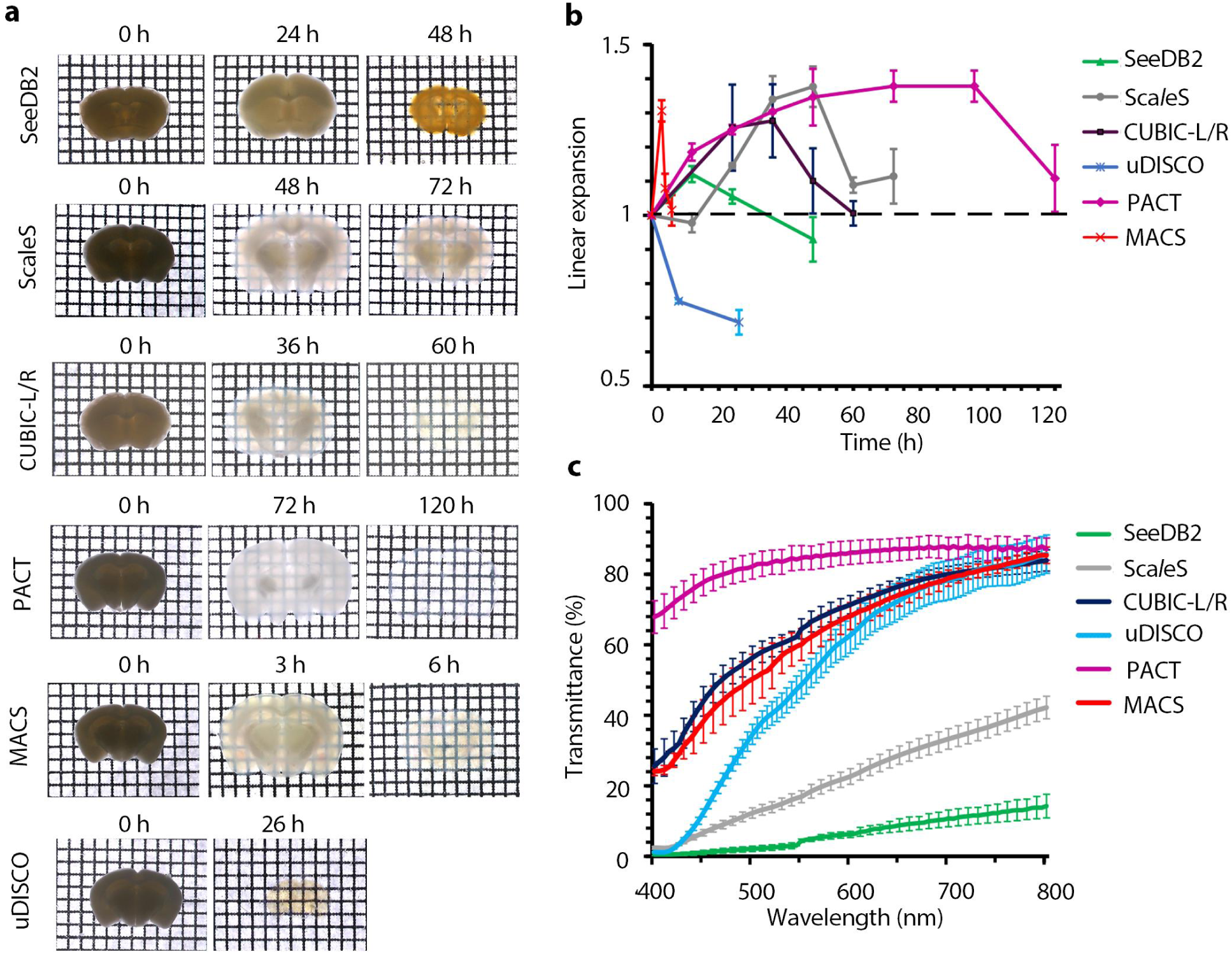
Clearing performance on 2-mm-thick brain slices for different methods. (**a**) Bright field images of samples before, during and after clearing for each clearing protocol. (**b**) Quantification of sample size changes in 2 mm adult slices (n=3) during different optical clearing procedures. (**c**) Transmission curves of 2 mm adult slices (n=3) cleared with various clearing protocols. All values are presented as the mean ± s.d.

**Fig. S2.**
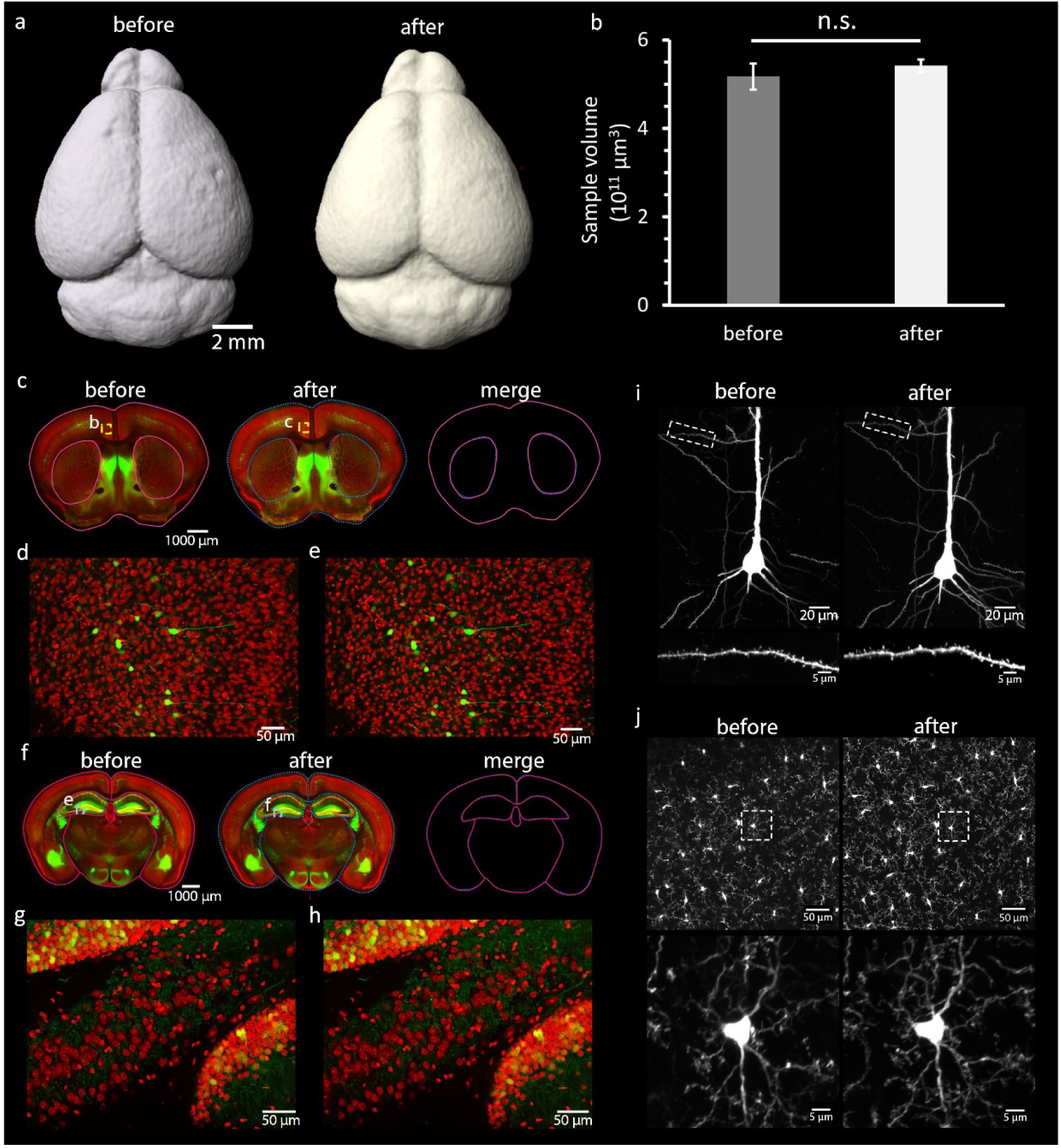
Preservation of tissue sizes and neural structures of brain samples after MACS treatment. (**a**) CT reconstruction of intact mouse brains before and after MACS clearing. (**b**) Measurement of volume change before and after MACS clearing. (**c**) Fluorescence images of coronal brain slices (1 mm, Bregma ≈ +0.50 mm) from adult *Thy1*-GFP-M mouse before and after MACS clearing. (**d, e**) Magnified images of boxed regions in (**c**). (**f**) Fluorescence images of coronal brain slices (1 mm, Bregma ≈ −1.70 mm) from adult *Thy1*-GFP-M mouse before and after MACS clearing. (**g, h**) Magnified images of boxed regions in (**f**). The borders of the main structures were traced and coloured magenta and blue, respectively. (**i, j**) Preservation of fine structures after MACS treatment. (**i**) Typical pyramidal neurons were imaged before and after MACS clearing. (**j**) Microglial cells were imaged before and after MACS clearing. Statistical significance in **b** (n.s., p>0.05) was assessed by an independent-sample t test (n=3).

**Fig. S3.**
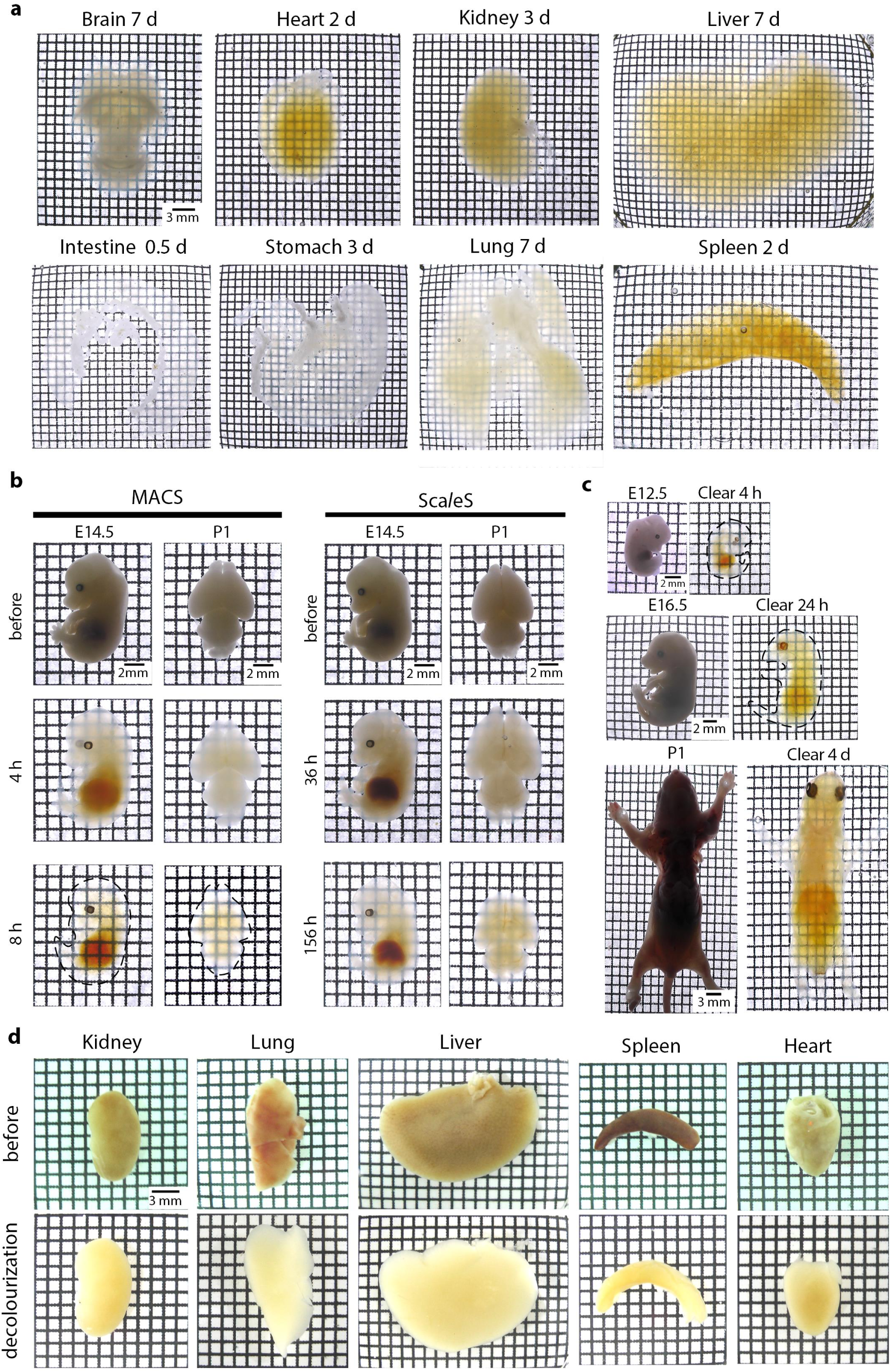
Efficient clearing and decolourization of different tissues by MACS. (**a**) Intact rat organs were cleared with MACS, and bright field images show that the brain, heart, kidney, liver, intestine, stomach, lung and spleen were rendered optically transparent at the indicated time. (**b**) Rapid clearing for mouse embryos and pups by MACS. Comparison of clearing performance for embryos and young mice by Sca*l*eS and MACS. E14.5 mouse embryos and whole neonatal mouse brain (P1) were used. Samples were cleared according to the original protocols for Sca*l*eS. At the time points indicated, bright field images were taken to examine the clarification of the samples. (**c**) Efficient clearing for embryos (E12.5 and E16.5) and P1 pups by MACS. (**d**) Decolourization of different mouse organs by MACS-R0. Hemi-rich mouse organs such as kidney, lung, liver, spleen and heart were incubated in MACS-R0 for 24 h and then washed with PBS. Reflection images were taken before and after treatment.

**Fig. S4.**
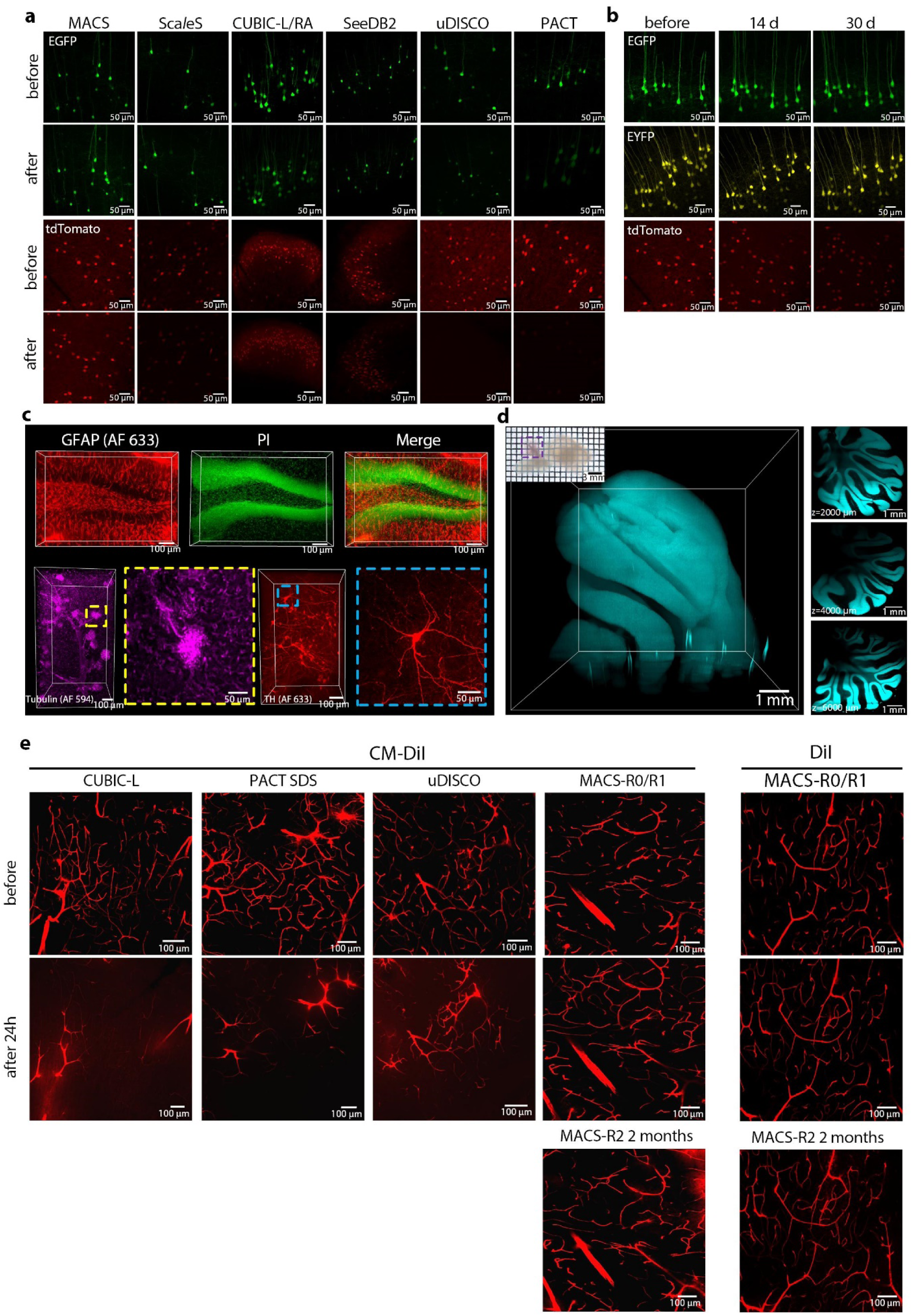
MACS performs good compatibility with transgenic labelling, immunolabelling, nucleus staining and DiI labelling, related to Fig. 2. (**a**) Fluorescence images of endogenous EGFP and tdTomato in 1 mm brain slices before and after clarification by each method. (**b**) Fluorescence images of endogenous EGFP, EYFP and tdTomato of MACS-cleared 1 mm brain slices over time. Bright field image of a rat hemisphere cleared by MACS. (**c**) (**top**) GFAP immunostaining and PI labelling of 1 mm coronal brain sections. Images show labelling of astrocytes and cell nuclei in hippocampus region. (**lower left**) 3D rendering of 1 mm mouse kidney slices stained by anti-beta tubulin, Magnification of boxed regions reveal fine labelling of microtubules. (**lower right**) 3D rendering of 1 mm brain blocks stained by anti-tyrosine hydrogenase (TH). Magnification of boxed regions reveals the structure of an individual TH-positive neuron. (**d**) LSFM imaging and reconstruction of the cerebellum of a PI-stained rat hemisphere cleared by MACS, the imaging depth is over 6800 μm. Optical sections are shown at different depths. The structure of the cerebellum is clearly visible throughout the entire imaging depth. (**e**) Comfocal images of CM-DiI labelled brain slices before and after 24 h clearing by each method. MACS could not only preserve the DiI signal, but also preserve the signals from CM-DiI.

**Fig. S5.**
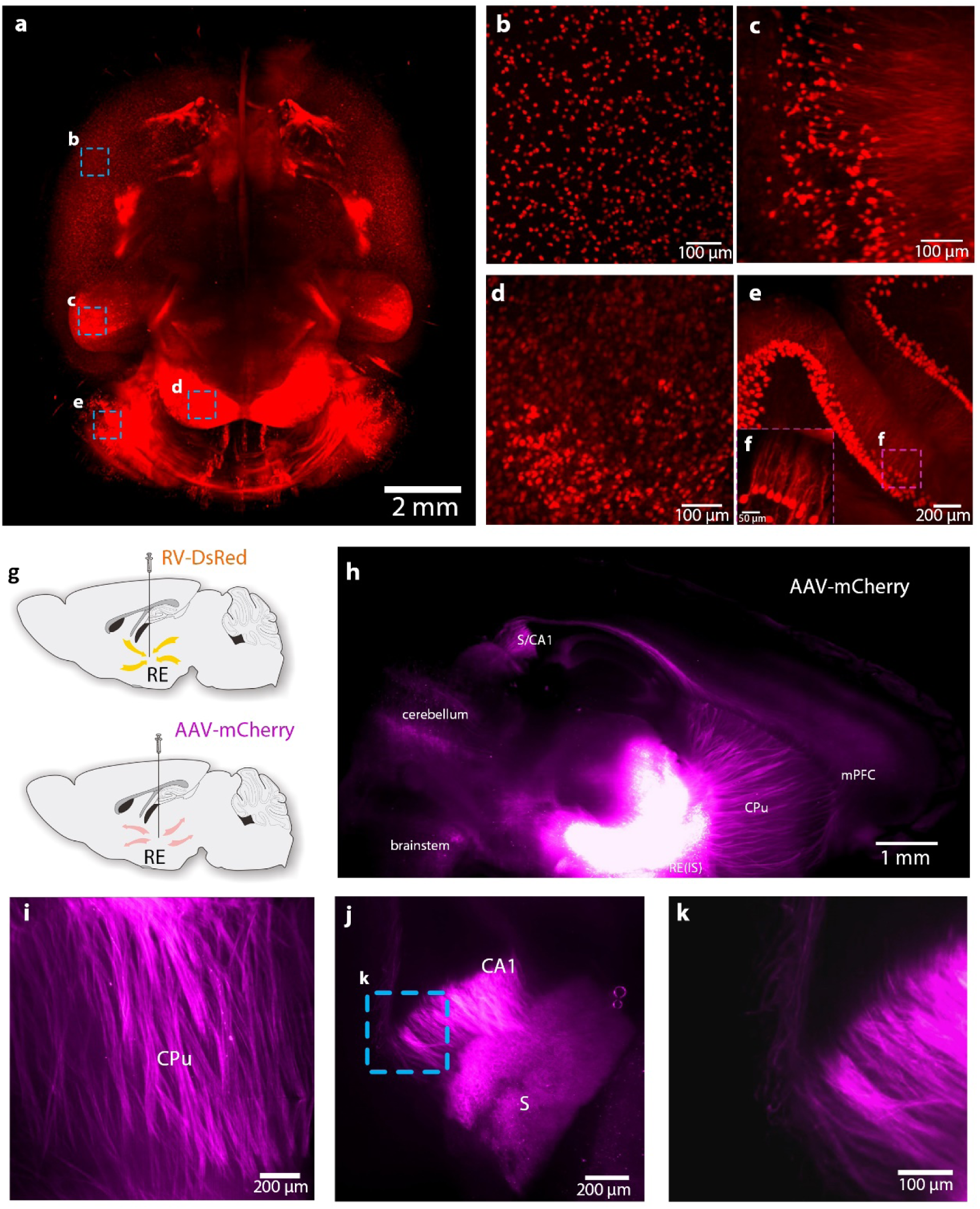
LSFM imaging of tdTomato-labelled and AAV-injected mouse brains cleared by MACS. (**a**) 3D reconstruction of LSFM images of a tdTomato-labelled (Sst-IRES-Cre::Ai14) whole brain cleared by MACS. (**b-e**) High-magnification images of tdTomato-positive neurons in the cortex (**b**), hippocampus (**c**), inferior colliculus (**d**) and cerebellum (**e**). (**f**) Details of the boxed region in **e**, showing the fine neuronal structures at single cell resolution. (**g**) Experimental design for RV and AAV injections. (**h**) 3D reconstruction of AAV-labelled projections from RE in the hemisphere. (**i**) Neural fibres projecting from RE to mPFC via CPu. (**j**) Projections to ventral CA1 of the hippocampus region and ventral subiculum (S). (**k**) The tiny fibres in d could be imaged. IS: injection site.

**Fig. S6.**
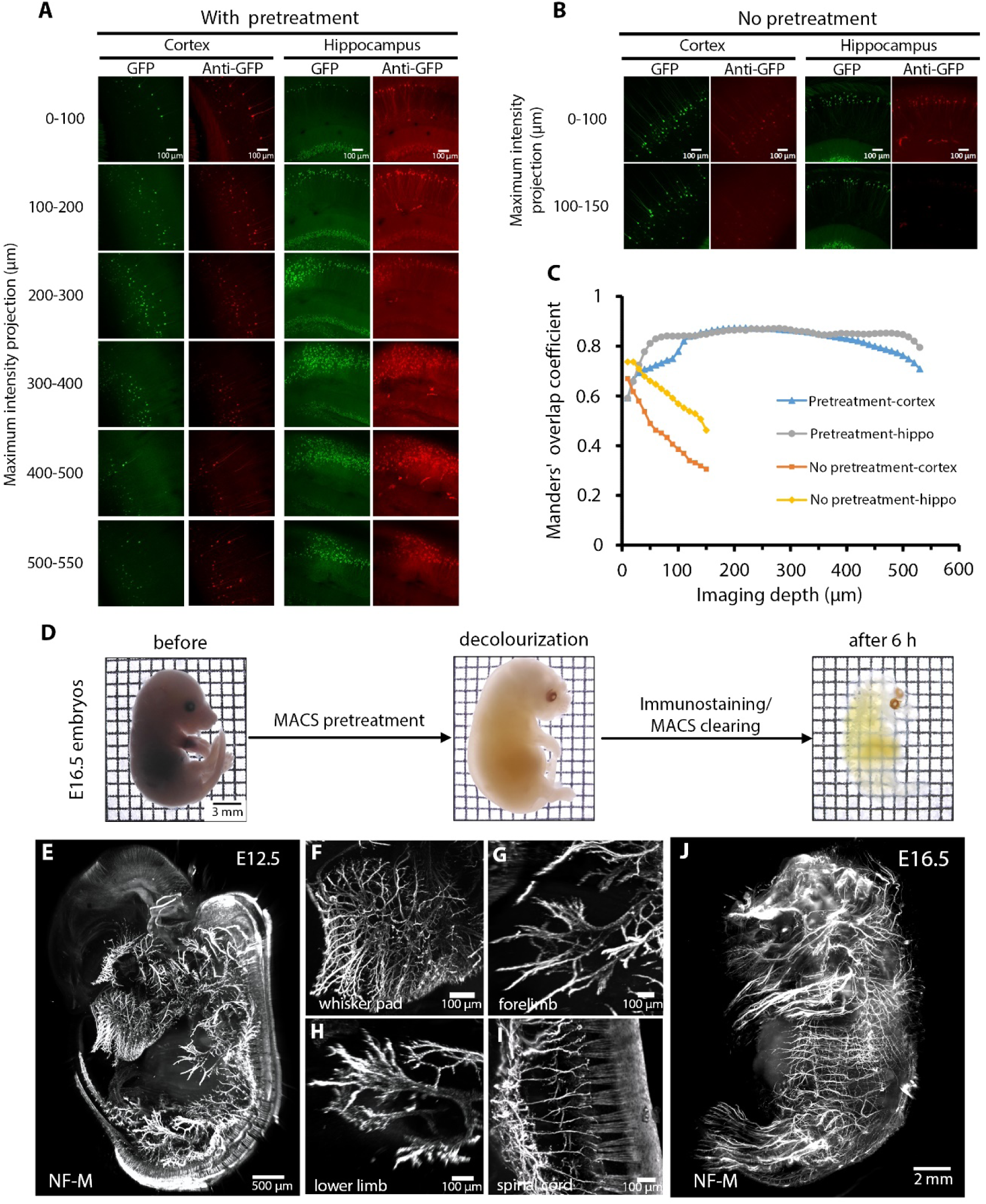
Whole-mount immunostaining of samples pre-treated by MXDA solution. (**a-b**) Promotion of antibody penetration of tissue treated by MACS solution. (**c**). Manders’ overlap coefficient was calculated by Fiji. (**d-j**) Whole-mount staining and imaging of intact embryos. (**d**) Work flow for immunostaining and clearing on whole embryos (e.g., E16.5). (**e**) Whole-mount immunostaining of E12.5 embryo labelled for neurofilament (NF-M). (**f-i**) Details of the labelled neural structures at E12.5 in the whisker pad (**f**), forelimb (**g**), lower limb (**h**) and spinal cord (**i**). (**j**) 3D rendering of whole E16.5 embryo labelled for neurofilament (NF-M).

**Fig. S7.**
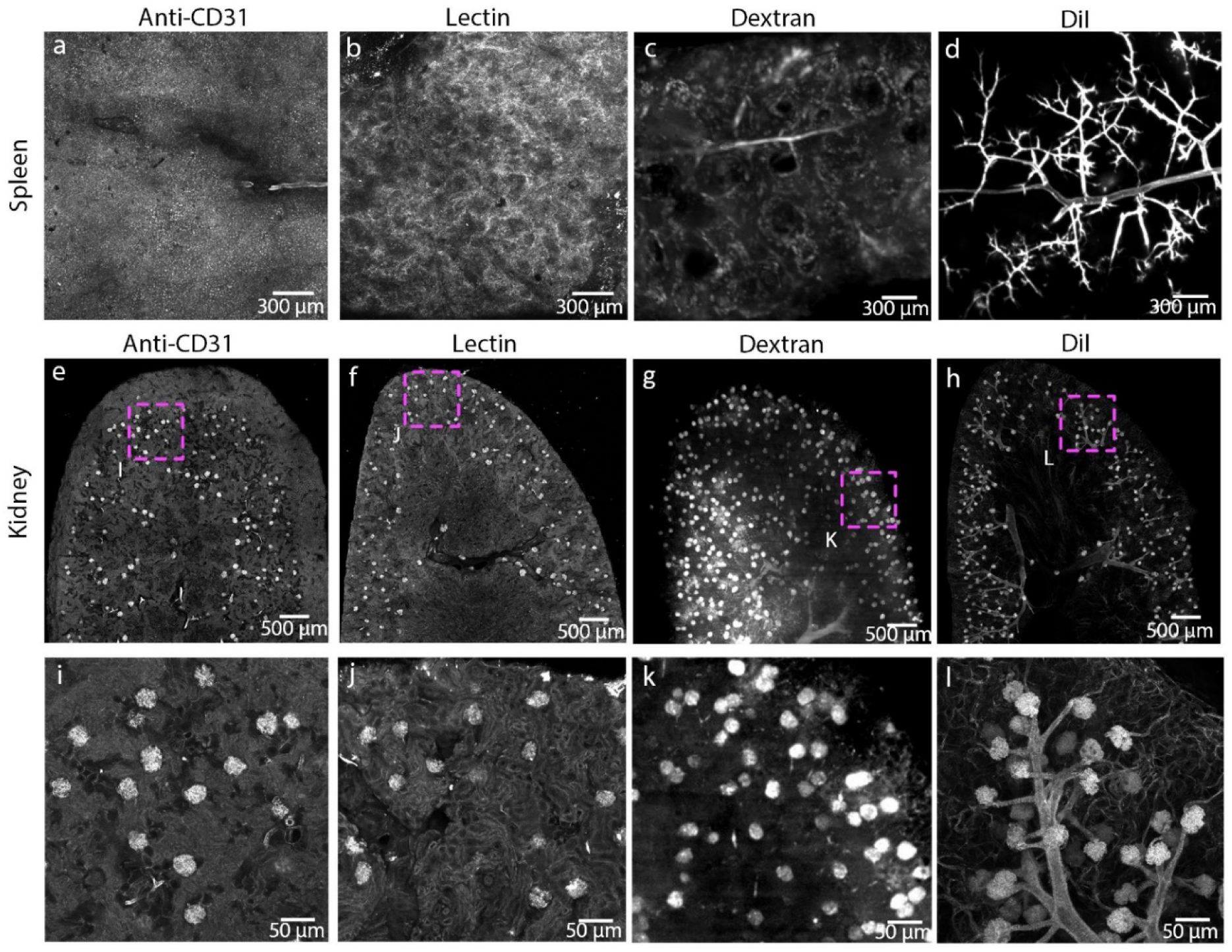
Comparison of different vasculature labelling methods on typical organs. (**a-d**) Vascular labelling of mouse spleens using anti-CD31 antibody (**a**), Lectin (**b**), Dextran (**c**) and DiI (**d**), respectively. (**e-h**) Vascular labelling of mouse kidneys using anti-CD31 antibody (**e**), lectin (**f**) dextran (**g**) and DiI (**h**), respectively. (**i-l**) Magnified images of boxed regions in **e-h**, respectively. The glomeruli were well labelled by three labelling methods, but the surrounding vessel structures were only seen in DiI labelled kidneys.

**Table S1.** Changes in the pH of different agents after decolourization.

| Agents | 50 vol% Quadrol | 0.01 M NaOH | 50 vol% MXDA |
| --- | --- | --- | --- |
| pH (before) | 10.5 | 12.0 | 11.5 |
| pH (after) | 9.8 | 10.6 | 11.4 |

The pH values of the three solutions were measured before and after decolourization of embryos containing blood. The pH values of Quadrol and NaOH solution revealed an obvious decline, while MXDA solution remained stable.

## References

1. Tomer, R. et al. SPED Light Sheet Microscopy: Fast Mapping of Biological System Structure and Function. Cell 163, 1796–1806 (2015).

2. Dodt, H. U. et al. Ultramicroscopy: three-dimensional visualization of neuronal networks in the whole mouse brain. Nat. Methods 4, 331 (2007).

3. Huisken, J. & Stainier, D.Y.R. Even fluorescence excitation by multidirectional selective plane illumination microscopy (mSPIM). Opt. Lett. 32, 2608–2610 (2007).

4. Livet, J. et al. Transgenic strategies for combinatorial expression of fluorescent proteins in the nervous system. Nature 450, 56–62 (2007).

5. Tuchin, V. V. et al. Light propagation in tissues with controlled optical properties. J. Biomed. Opt. 2, 401–417 (1997).

6. Economo, M.N. et al. A platform for brain-wide imaging and reconstruction of individual neurons. eLife 5, e10566 (2016).

7. Gong, H. et al. Continuously tracing brain-wide long-distance axonal projections in mice at a one-micron voxel resolution. NeuroImage 74, 87–98 (2013).

8. Li, A. et al. Micro-Optical Sectioning Tomography to Obtain a High-Resolution Atlas of the Mouse Brain. Science 330, 1404 (2010).

9. Seiriki, K. et al. High-Speed and Scalable Whole-Brain Imaging in Rodents and Primates. Neuron 94, 1085–1100 e1086 (2017).

10. Susaki, E.A. & Ueda, H.R. Whole-body and Whole-Organ Clearing and Imaging Techniques with Single Cell Resolution: Toward Organism-Level Systems Biology in Mammals. Cell Chem. Biol. 23, 137–157 (2016).

11. Tainaka, K., Kuno, A., Kubota, S.I., Murakami, T. & Ueda, H.R. Chemical Principles in Tissue Clearing and Staining Protocols for Whole-Body Cell Profiling. Annu. Rev. Cell Dev. Biol. 32, 713–741 (2016).

12. Zhu, D., Larin, K.V., Luo, Q. & Tuchin, V.V. Recent progress in tissue optical clearing. Laser Photon. Rev. 7, 732–757 (2013).

13. Yu, T., Qi, Y., Gong, H., Luo, Q. & Zhu, D. Optical clearing for multiscale biological tissues. J. Biophotonics 11 (2018).

14. Erturk, A. et al. Three-dimensional imaging of solvent-cleared organs using 3DISCO. Nat. Protoc. 7, 1983–1995 (2012).

15. Erturk, A. et al. Three-dimensional imaging of the unsectioned adult spinal cord to assess axon regeneration and glial responses after injury. Nat. Med. 18, 166–171 (2011).

16. Renier, N. et al. iDISCO: a simple, rapid method to immunolabel large tissue samples for volume imaging. Cell 159, 896–910 (2014).

17. Pan, C. et al. Shrinkage-mediated imaging of entire organs and organisms using uDISCO. Nat. Methods 13, 859–867 (2016).

18. Jing, D. et al. Tissue clearing of both hard and soft tissue organs with the PEGASOS method. Cell Res. 28, 803–818 (2018).

19. Klingberg, A. et al. Fully Automated Evaluation of Total Glomerular Number and Capillary Tuft Size in Nephritic Kidneys Using Lightsheet Microscopy. J. Am. Soc. Nephrol. 28, 452–459 (2017).

20. Qi, Y. et al. FDISCO: Advanced solvent-based clearing method for imaging whole organs. Sci. Adv. 5, eaau8355 (2019).

21. Cai, R. et al. Panoptic imaging of transparent mice reveals whole-body neuronal projections and skull-meninges connections. Nat. Neurosci. 22, 317–327 (2019).

22. Hahn, C. et al. High-resolution imaging of fluorescent whole mouse brains using stabilised organic media (sDISCO). J.Biophotonics, e201800368 (2019).

23. Wang, Q., Liu, K., Yang, L., Wang, H. & Yang, J. BoneClear: whole-tissue immunolabeling of the intact mouse bones for 3D imaging of neural anatomy and pathology. Cell Res. (2019).

24. Ke, M.T. et al. Super-Resolution Mapping of Neuronal Circuitry With an Index-Optimized Clearing Agent. Cell Rep. 14, 2718–2732 (2016).

25. Ke, M.T., Fujimoto, S. & Imai, T. SeeDB: a simple and morphology-preserving optical clearing agent for neuronal circuit reconstruction. Nat. Neurosci. 16, 1154–1161 (2013).

26. Kuwajima, T. et al. ClearT: a detergent- and solvent-free clearing method for neuronal and non-neuronal tissue. Development 140, 1364–1368 (2013).

27. Hama, H. et al. Sca*l*eS: an optical clearing palette for biological imaging. Nat. Neurosci. 18, 1518–1529 (2015).

28. Susaki, Etsuo A. et al. Whole-Brain Imaging with Single-Cell Resolution Using Chemical Cocktails and Computational Analysis. Cell 157, 726–739 (2014).

29. Tainaka, K. et al. Whole-body imaging with single-cell resolution by tissue decolorization. Cell 159, 911–924 (2014).

30. Kubota, S.I. et al. Whole-Body Profiling of Cancer Metastasis with Single-Cell Resolution. Cell Rep. 20, 236–250 (2017).

31. Tainaka, K. et al. Chemical Landscape for Tissue Clearing Based on Hydrophilic Reagents. Cell Rep. 24, 2196–2210 e2199 (2018). 21, 625-637 (2018).

32. Chung, K. et al. Structural and molecular interrogation of intact biological systems. Nature 497, 332–337 (2013).

33. Tomer, R., Ye, L., Hsueh, B. & Deisseroth, K. Advanced CLARITY for rapid and high-resolution imaging of intact tissues. Nat. Protoc. 9, 1682–1697 (2014).

34. Yang, B. et al. Single-Cell Phenotyping within Transparent Intact Tissue through Whole-Body Clearing. Cell 158, 945–958 (2014).

35. Murray, E., et al. Simple, Scalable Proteomic Imaging for High-Dimensional Profiling of Intact Systems. Cell 163, 1500–1514 (2015).

36. Lai, H.M. et al. Next generation histology methods for three-dimensional imaging of fresh and archival human brain tissues. Nat. Commun. 9, 1066 (2018).

37. Li, W., Germain, R.N. & Gerner, M.Y. Multiplex, quantitative cellular analysis in large tissue volumes with clearing-enhanced 3D microscopy (Ce3D). Proc. Natl. Acad. Sci. U.S.A. 114, E7321–E7330 (2017).

38. Yu, T. et al. RTF: a rapid and versatile tissue optical clearing method. Sci. Rep. 8, 1964 (2018).

39. Wan, P. et al. Evaluation of seven optical clearing methods in mouse brain. Neurophotonics 5, 035007 (2018).

40. Collazo, A., Bronner-Fraser, M. & Fraser, S.E. Vital dye labelling of Xenopus laevis trunk neural crest reveals multipotency and novel pathways of migration. Development 118, 363 (1993).

41. Stark, M.R., Sechrist, J., Bronner-Fraser, M. & Marcelle, C. Neural tube-ectoderm interactions are required for trigeminal placode formation. Development 124, 4287 (1997).

42. Molnár, Z. & Blakemore, C. Lack of regional specificity for connections formed between thalamus and cortex in coculture. Nature 351, 475 (1991).

43. Jensen, K.H. & Berg, R.W. CLARITY-compatible lipophilic dyes for electrode marking and neuronal tracing. Sci. Rep. 6, 32674 (2016).

44. Kim, J.H. et al. Optimizing tissue-clearing conditions based on analysis of the critical factors affecting tissue-clearing procedures. Sci. Rep. 8, 12815 (2018).

45. Wickersham, I.R., Finke, S., Conzelmann, K.K. & Callaway, E.M. Retrograde neuronal tracing with a deletion-mutant rabies virus. Nat. Methods 4, 47–49 (2007).

46. Nassi, J.J., Cepko, C.L., Born, R.T. & Beier, K.T. Neuroanatomy goes viral! *Front*. Neuroanat. 9, 80 (2015).

47. Griffin, A.L. Role of the thalamic nucleus reuniens in mediating interactions between the hippocampus and medial prefrontal cortex during spatial working memory. Front. Syst. Neurosci. 9, 29 (2015).

48. McKenna, J.T. & Vertes, R.P. Afferent projections to nucleus reuniens of the thalamus. J. Comp. Neurol. 480, 115–142 (2004).

49. Mj, D.V.D.W. & Witter, M.P. Projections from the nucleus reuniens thalami to the entorhinal cortex, hippocampal field CA1, and the subiculum in the rat arise from different populations of neurons. J Comp Neurol. 364, 637–650 (2015).

50. Xiong, B. et al. Precise Cerebral Vascular Atlas in Stereotaxic Coordinates of Whole Mouse Brain, Front. Neuroanat. 11, 128 (2017).

51. Zhang, L.Y. et al. CLARITY for High-resolution Imaging and Quantification of Vasculature in the Whole Mouse Brain. Aging and disease 9, 262–272 (2018).

52. Li, Y. et al. Direct labeling and visualization of blood vessels with lipophilic carbocyanine dye DiI. Nat. Protoc. 3, 1703–1708 (2008).

53. Kim, S. Y. et al. Stochastic electrotransport selectively enhances the transport of highly electromobile molecules. Proc. Natl. Acad. Sci. U.S.A. 112, E6274 (2015).

54. Lee, E. et al. ACT-PRESTO: Rapid and consistent tissue clearing and labeling method for 3-dimensional (3D) imaging. Sci. Rep. 6, 18631 (2016).

55. Faber, D.J., Mik, E.G., Aalders, M.C.G. & Leeuwen, T.G., Van Light absorption of (oxy-)hemoglobin assessed by spectroscopic optical coherence tomography. Opt. Lett. 28, 1436–1438 (2003).

56. Yang, X. et al. Combined system of fluorescence diffuse optical tomography and microcomputed tomography for small animal imaging. Rev. Sci. Instrum. 81, 054304 (2010).

